## Supplementary Figures and Notes for "A complex between IF2 and NusA links transcription and translation"

#### Impact of HaloTag fusion on the functionality of initiation factors

To explored if the addition of a HaloTag compromises IF activity we constructed plasmids for IPTG-inducible expression of individual non-labeled IFs, as well as N-terminal HaloTag fusions of each IF. To test whether there is any growth defect associated with the addition of the HaloTag, the plasmids were introduced in *E. coli* strains in which the corresponding IFs were deleted from the genome (Supplementary Fig. 1). Without IF induction, all strains showed poor growth, suggesting that leakage level expression is insufficient. When IPTG was present in the media at 32-64 µM, the strains with plasmid-expressed IF2α, IF3, and IF1 showed doubling times similar to that of the wild-type (WT) strain carrying an empty plasmid, suggesting that at this induction level, the plasmids support fast growth (Supplementary Fig. 1a,b,d). Higher expression of IF3 inhibited growth, suggesting that proper cell physiology requires a fine-tuned concentration of this factor, in line with in vitro results showing inhibition of translation initiation at high IF3 levels (Supplementary Fig. 1d) [1]. Expression of the IF2γ isoform also supported growth, albeit at a slower rate even at higher concentrations of the inducer (Supplementary Fig. 1c).

For neither of the IF2 isoforms, the addition of HaloTag at the N-terminus altered the growth rate, indicating that HaloTag does not interfere with the factors' functions (Supplementary Fig. 1b,c). Induction of the HaloTag-IF3 fusion could also support fast growth of the strain lacking chromosomal IF3, albeit at a slightly higher concentration of the inducer than needed for the non-HaloTag version (Supplementary Fig. 1d). This indicates that the addition of the HaloTag might have a slight negative effect on IF3 function, but the discrepancy could also be due to changes in expression level or folding efficiency because of the HaloTag addition. Finally, the strain lacking IF1 and expressing HaloTag-IF1 shows noticeable growth defect in comparison with the strain where WT IF1 is expressed from a plasmid at any level of induction, showing that the HaloTag negatively affects the functionality of IF1 (Supplementary Fig. 1a).

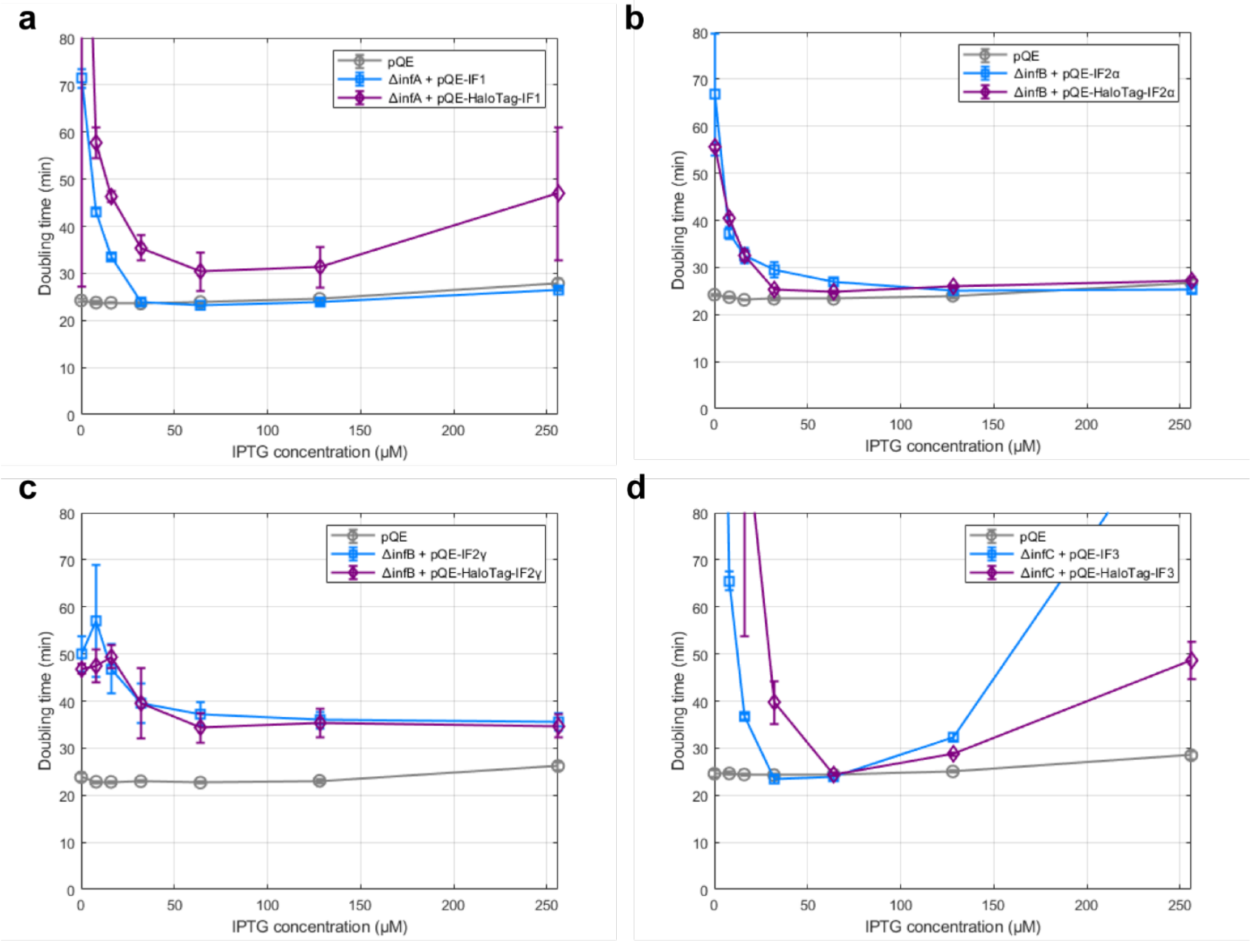


**Supplementary Fig. 1. Effect of N-terminal HaloTag fusion on activity of IFs. a.** Effect of IPTG concentration on doubling time of *E. coli* strain carrying an empty pQE plasmid and *E. coli ΔinfA* strain carrying either a plasmid pQE-IF1 expressing IF1 protein or a plasmid pQE-HaloTag-IF1 expressing HaloTag-IF1 fusion. **b.** Effect of IPTG concentration on doubling time of *E. coli* strain carrying an empty pQE plasmid and *E. coli ΔinfB* strain carrying either a plasmid pQE-IF2α expressing α isoform of IF2 protein or a plasmid pQE-HaloTag-IF2α expressing HaloTag-IF2α fusion protein. **c.** Effect of IPTG concentration on doubling time of *E. coli* strain carrying an empty pQE plasmid and *E. coli ΔinfB* strain carrying either a plasmid pQE-IF2γ expressing γ isoform of IF2 protein or a plasmid pQE-HaloTag-IF2γ expressing HaloTag-IF2γ fusion protein. **d.** Effect of IPTG concentration on doubling time of *E. coli* strain carrying an empty pQE plasmid and *E. coli ΔinfC* strain carrying either a plasmid pQE-IF3 expressing IF3 protein or a plasmid pQE-HaloTag-IF3 expressing HaloTag-IF3 fusion.

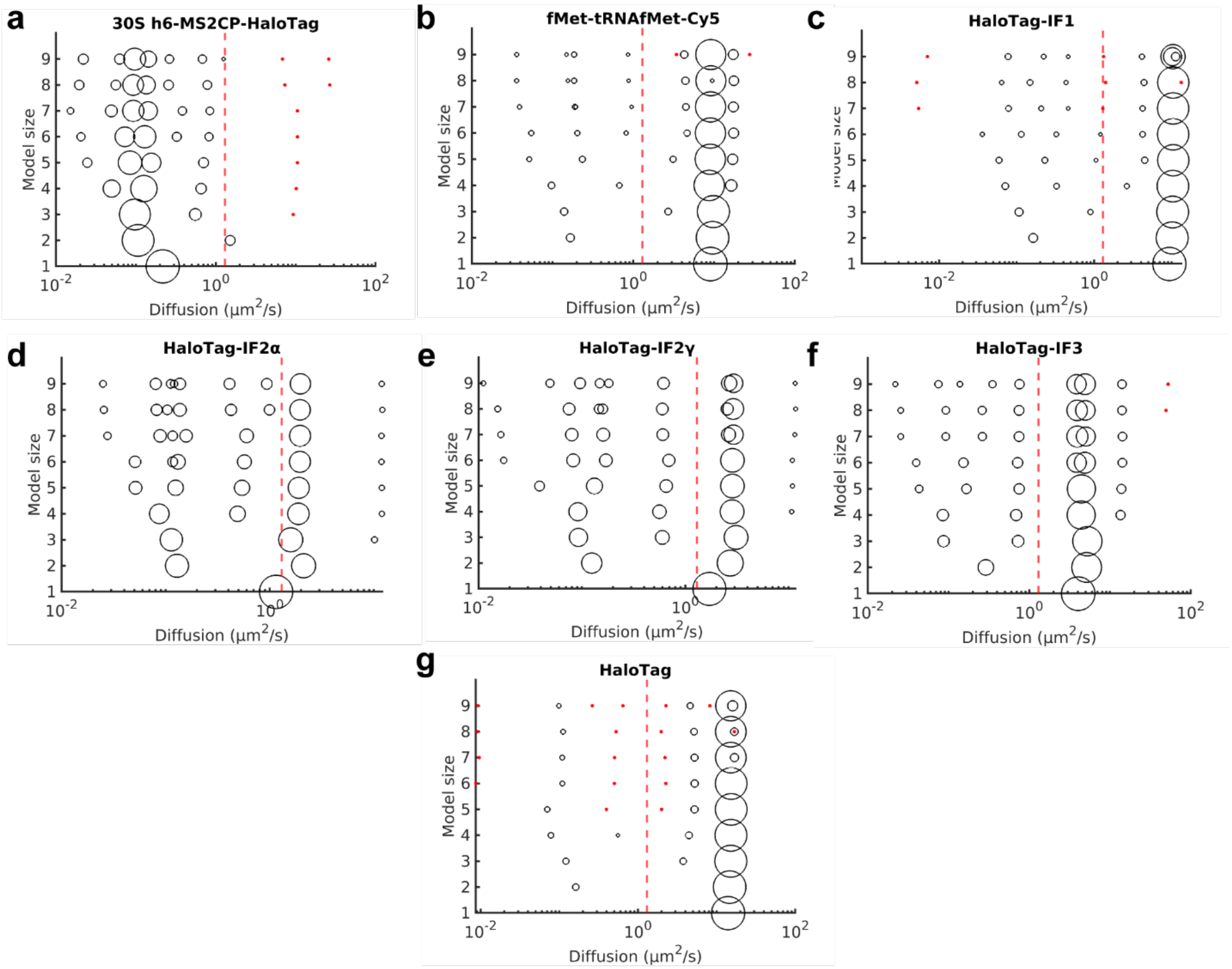


**Supplementary Fig. 2. Fitted HMM models of diffusional states for 30S, HaloTag, fMet-tRNA$^{fMet}$, and all labeled IFs.** The area of the circles represents the relative occupancy in different diffusional states. Diffusional states with occupancies lower than 1% are marked as *. The red dashed line marks the threshold at 1.3 μm$^2$/s corresponding to the upper limit of diffusional states detected for 30S. All models are shown in Supplementary Data 1-7.

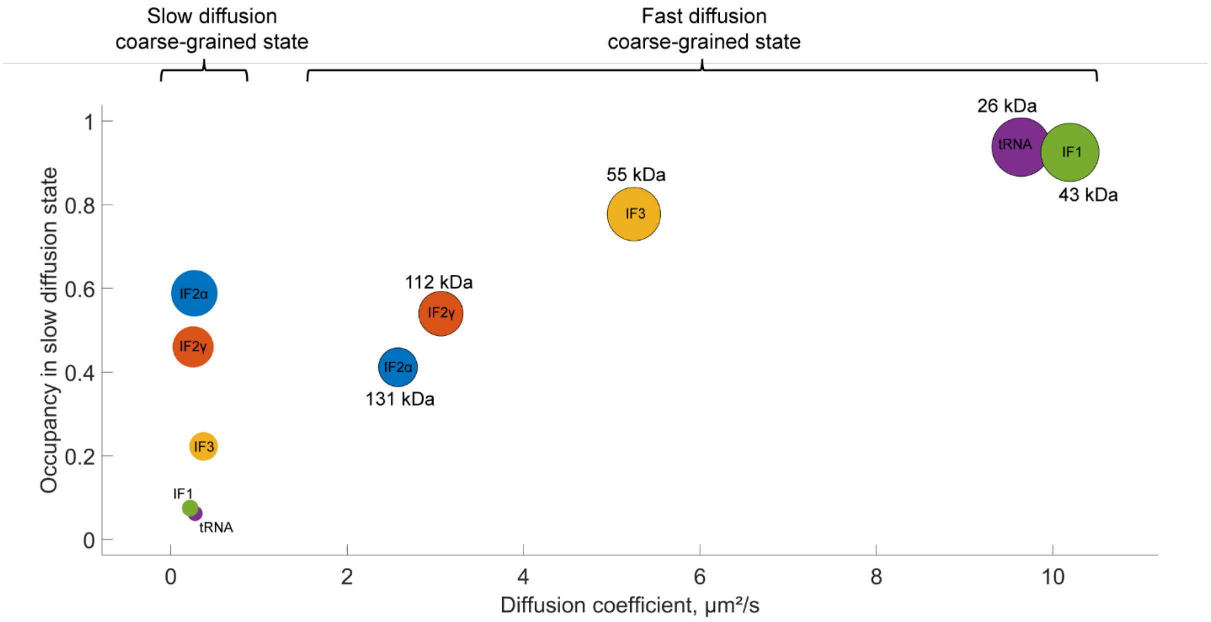


**Supplementary Fig. 3. Coarse-grained results of HMM modelling.** Estimated steady-state occupancies of HaloTag-IF2α, HaloTag-IF2γ, HaloTag-IF3, and fMet-[Cy5]tRNA$^{fMet}$ in the slow and fast diffusion states after coarse-graining from HMM-fitted model sizes 6–9 (Supplementary Data 8). The area of the circles represents the relative occupancy in a coarse-grained diffusional state.

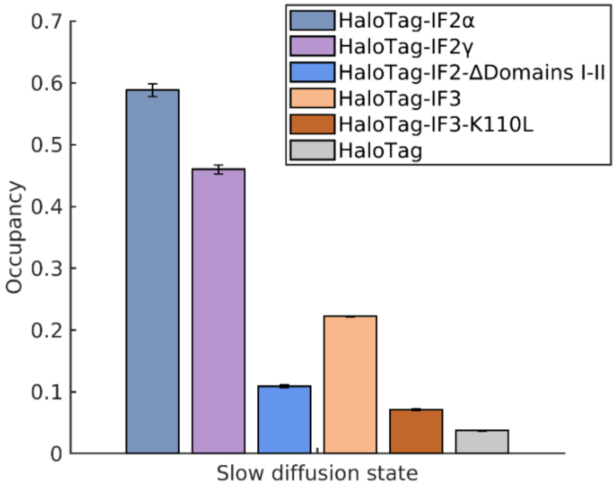


**Supplementary Fig. 4. Effect of mutation on slow diffusion state occupancy of HaloTag-IF2 and HaloTag-IF3.** Estimated steady-state occupancies of HaloTag-IF2α, HaloTag-IF2γ, variant of HaloTag-IF2 lacking Domain I and II, HaloTag-IF3, HaloTag-IF3-K110L mutant, and HaloTag in the slow diffusion state. The data shows weighted averages from coarse-grained HMM-fitted model sizes 6–9 (Supplementary Data 8). Results for individual model sizes (1–9) are shown in Supplementary Data 1-3, 9-10. Error bars represent weighted standard deviation, calculated from individual model sizes (6–9).

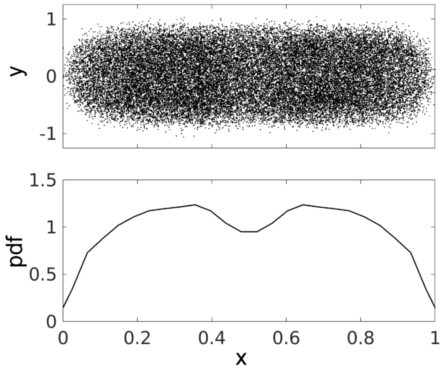


**Supplementary Fig. 5**. **Spatial distribution of HaloTag-IF2-Domain-I in the fast diffusional state from a 2-state HMM model.** In the top panel, dot locations are plotted on normalized cell coordinates (x = long cell axis, and y = short cell axis). The bottom panel shows the distribution of dot coordinates projected on the long cell axis (only middle 40% of the cells are included in the plot to avoid possible artifacts due to proximity to the membrane).

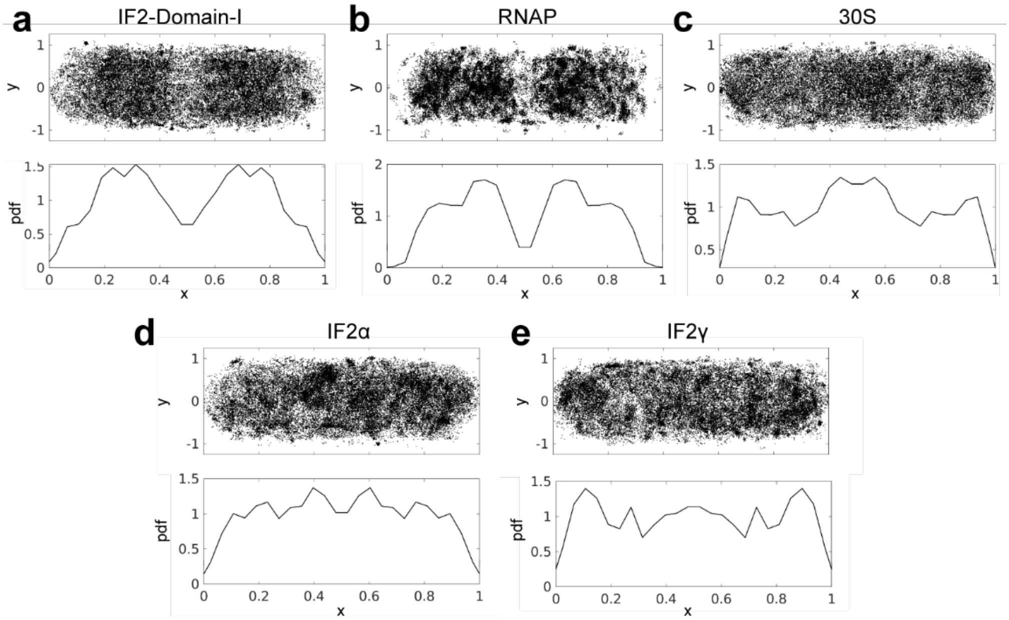


**Supplementary Fig. 6 (related to Fig. 3). Spatial distribution of the slow diffusional state for 30S, RNAP, and IF2 variants.** Spatial distribution of HaloTag-IF2-Domain-I (**a**), RpoC-HaloTag (**b**), h6-MS2CP-HaloTag labeled 30S (**c**), HaloTag-IF2α (**d**), HaloTag-IF2γ (**e**) in the slow diffusional state from 2-state HMM models. In the top panels, dot locations are plotted on normalized cell coordinates (x = long cell axis, and y = short cell axis). The bottom panels show the distribution of dot coordinates projected on the long cell axis (only middle 40% of the cells are included in the plot to avoid possible artifacts due to proximity to the membrane).

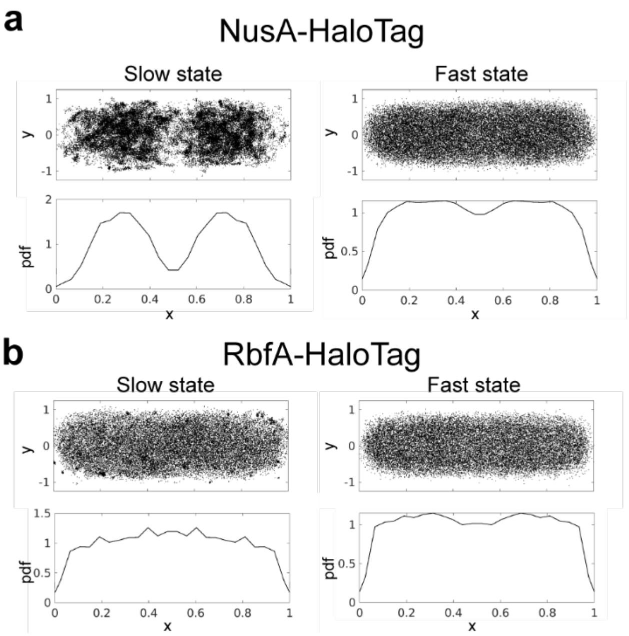


**Supplementary Fig. 7 (related to Fig. 4). Spatial distribution of NusA-HaloTag and RbfA-HaloTag in the fast and slow diffusional state from 2-state HMM models.** In the top panel, dot locations are plotted on normalized cell coordinates (x = long cell axis, and y = short cell axis). The bottom panels show the distribution of dot coordinates projected on the long cell axis (only middle 40% of the cells are included in the plot to avoid possible artifacts due to proximity to the membrane).

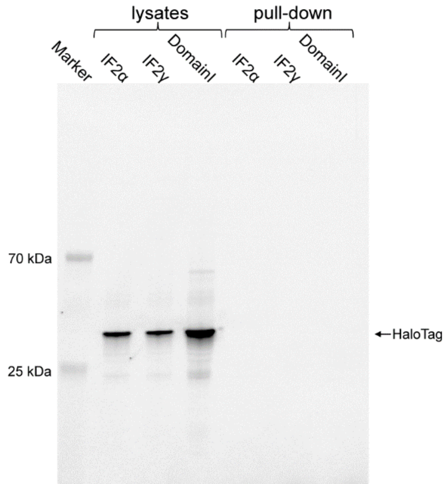


**Supplementary Fig. 8. SDS-PAGE analysis of cell lysates and affinity purification fractions for the presence of HaloTag.** 6His-IF2α, 6His-IF2γ, and IF2-Domain-I-6His were overexpressed in an *E. coli* strain carrying a plasmid for expression of HaloTag. Lysates and samples affinity purified on Co-NTA agarose (i.e. "pull-down") were loaded on SDS-PAGE gel and stained using the JFX549 dye to label the HaloTag.

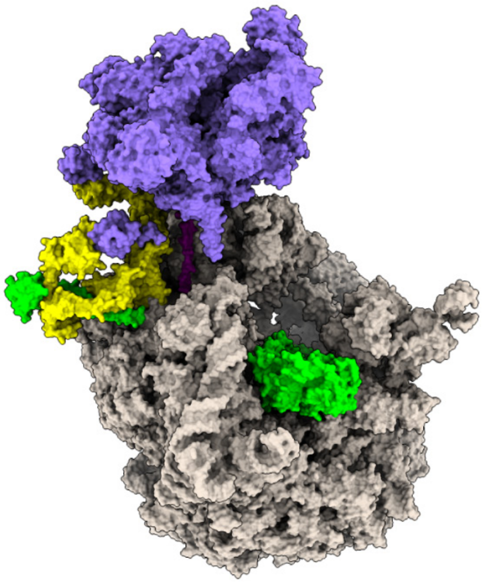

**Supplementary Fig. 9. Modelling of a coupled transcription-translation complex with fitted IF2.** Structure of a coupled transcription-translation complex containing NusA and NusG (6X7F PDB) was aligned with the structure of ribosome-bound IF2 (3JCJ PDB) to model the position of the C-terminal part of IF2 (amino acids 382-890) and aligned with the Alphafold-modeled complex between NusA and N-terminal fragment of IF2 (amino acids 2-94). Both fitted fragments of IF2 are shown in green, NusA is shown in yellow, RNAP is shown in purple, ribosome is shown in grey.

### Supplementary Note 1.

Elongation rate of the ribosome growing with the growth rate at 2 doublings/h (matching the growth rate of the strains in microscopy experiments) has been estimated to be 16-17 amino acids per second in studies where direct measurements of the elongation rate were performed [2]. Indirect calculations based on the amount of total protein per cell, the number of ribosomes per cell, the percentage of active ribosomes (set at 85%), and the cell growth rate, provide an estimate for the global elongation rate at 21-22 amino acid per second [3].

Estimates for the average length of a “typical” E. coli protein varies between different approaches, where ribosome profiling data provide an estimate at 192-222 amino acids [4] (see Supplementary Data 15 in [5]), and estimation based on proteomics data provides an estimate at 242 amino acids [6] (see Supplementary Data 14 in [5]). Taking these estimates into account elongation of a ‘typical’ E. coli protein by the ribosome requires ≈9-15 seconds. Our previous results for tracking of the ribosomal subunits provide an estimate for an average time required for initiation for a ribosome at 1.3-1.6 s, i.e. time required for initiation. Assuming that the termination of translation proceeds fast, we estimate that the average cycle time of the ribosome to translate a typical protein lies between 10-17 s.
